## Supplementary figures and images for "Alpha-Band Phase Modulates Perceptual Sensitivity by Changing Internal Noise and Sensory Tuning"

### Supplemental Figure 1

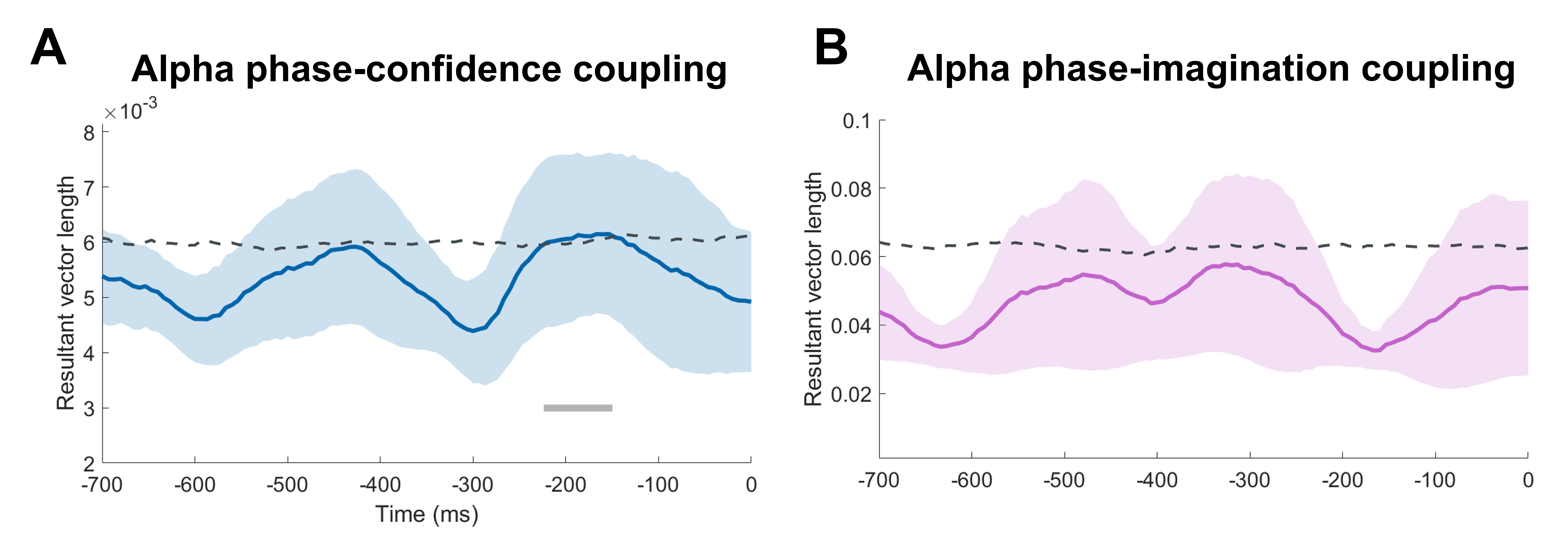

### Supplemental Figure 2

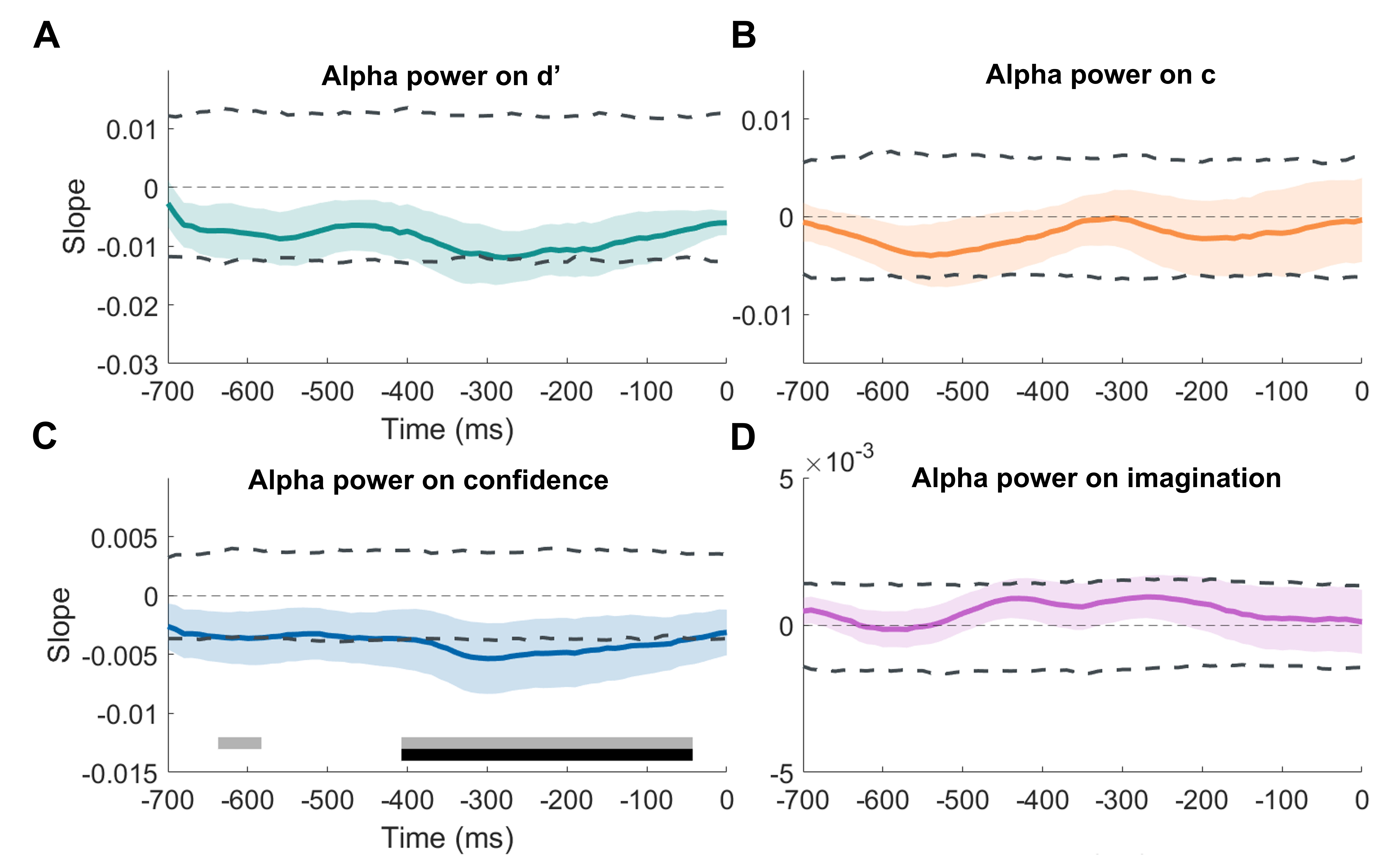

### Supplemental Figure 3

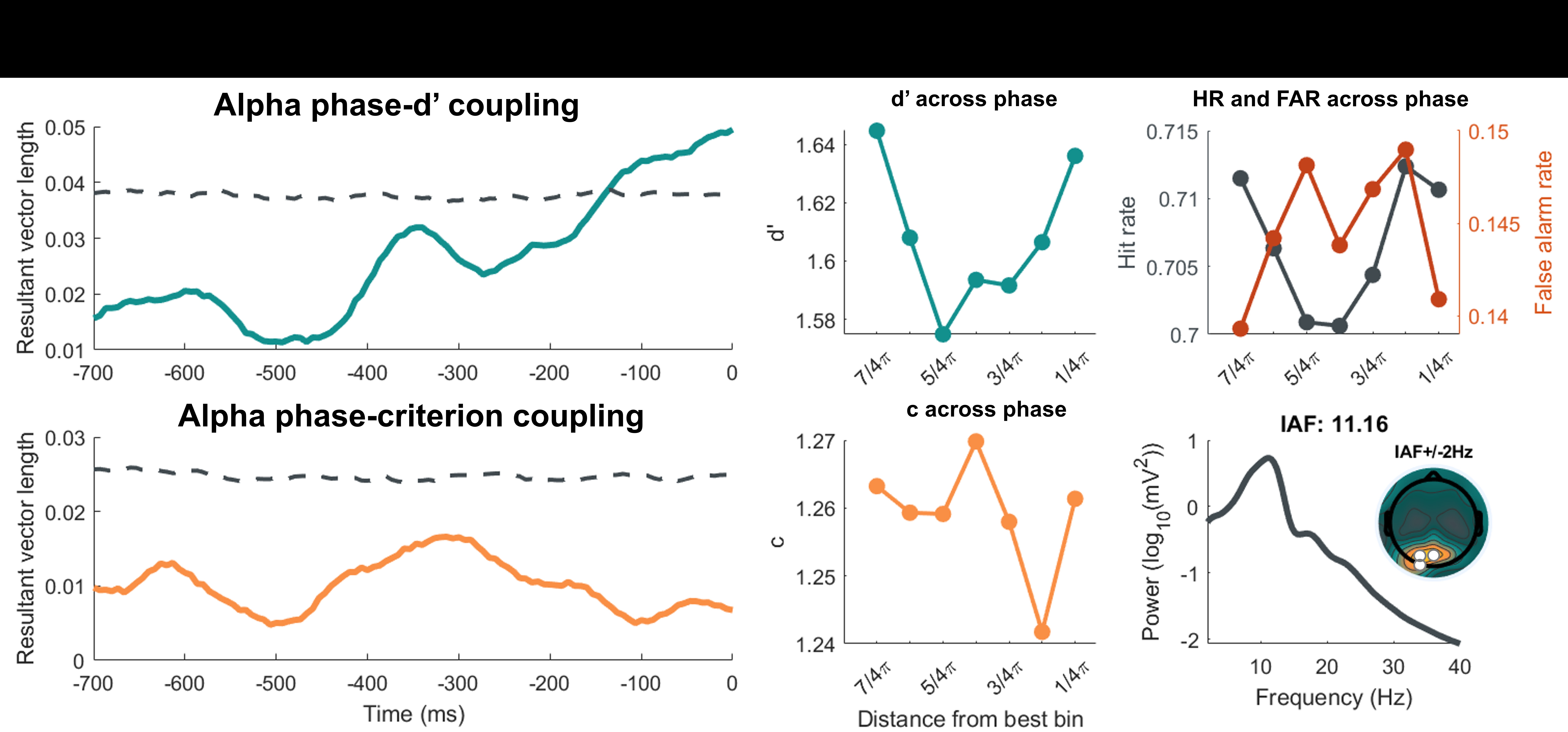

### Supplemental Figure 3

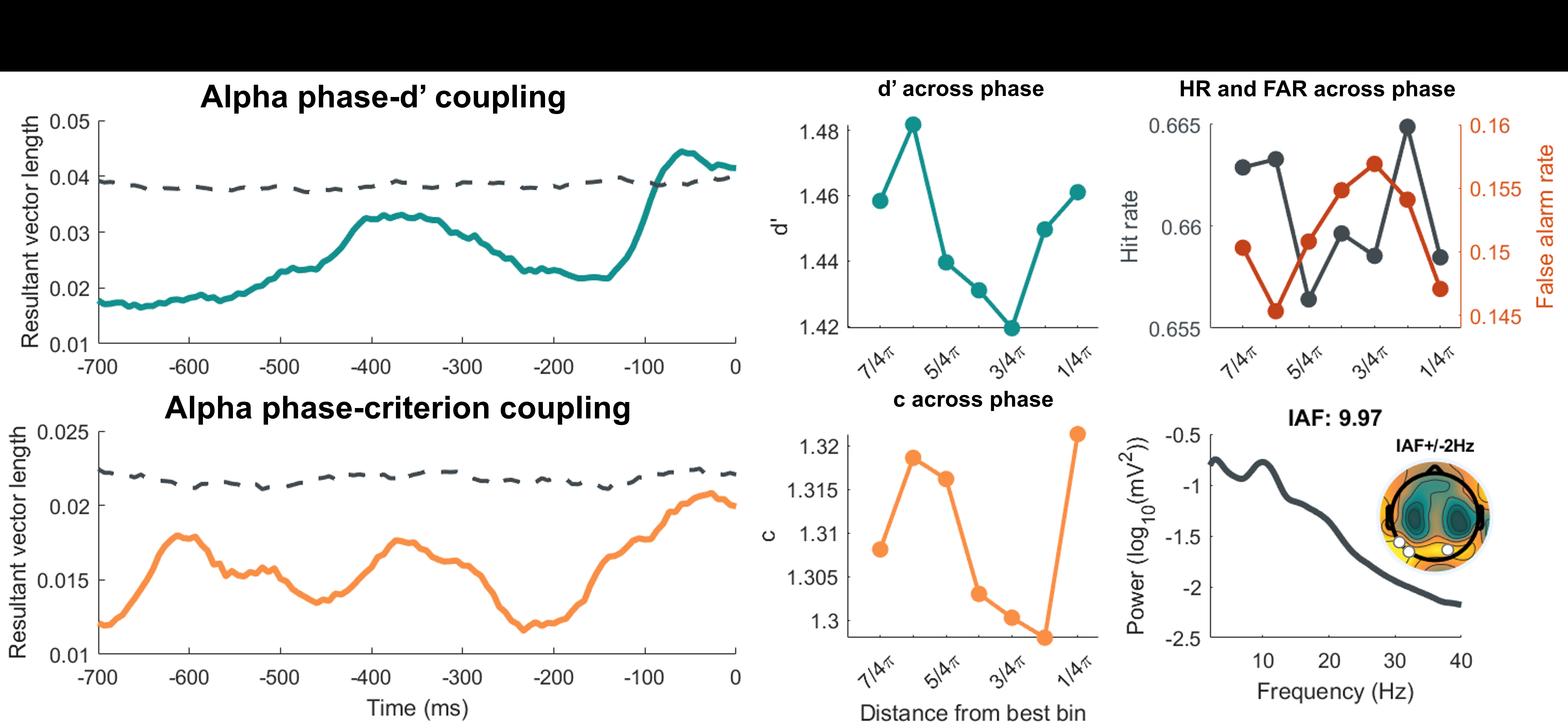

### Supplemental Figure 3

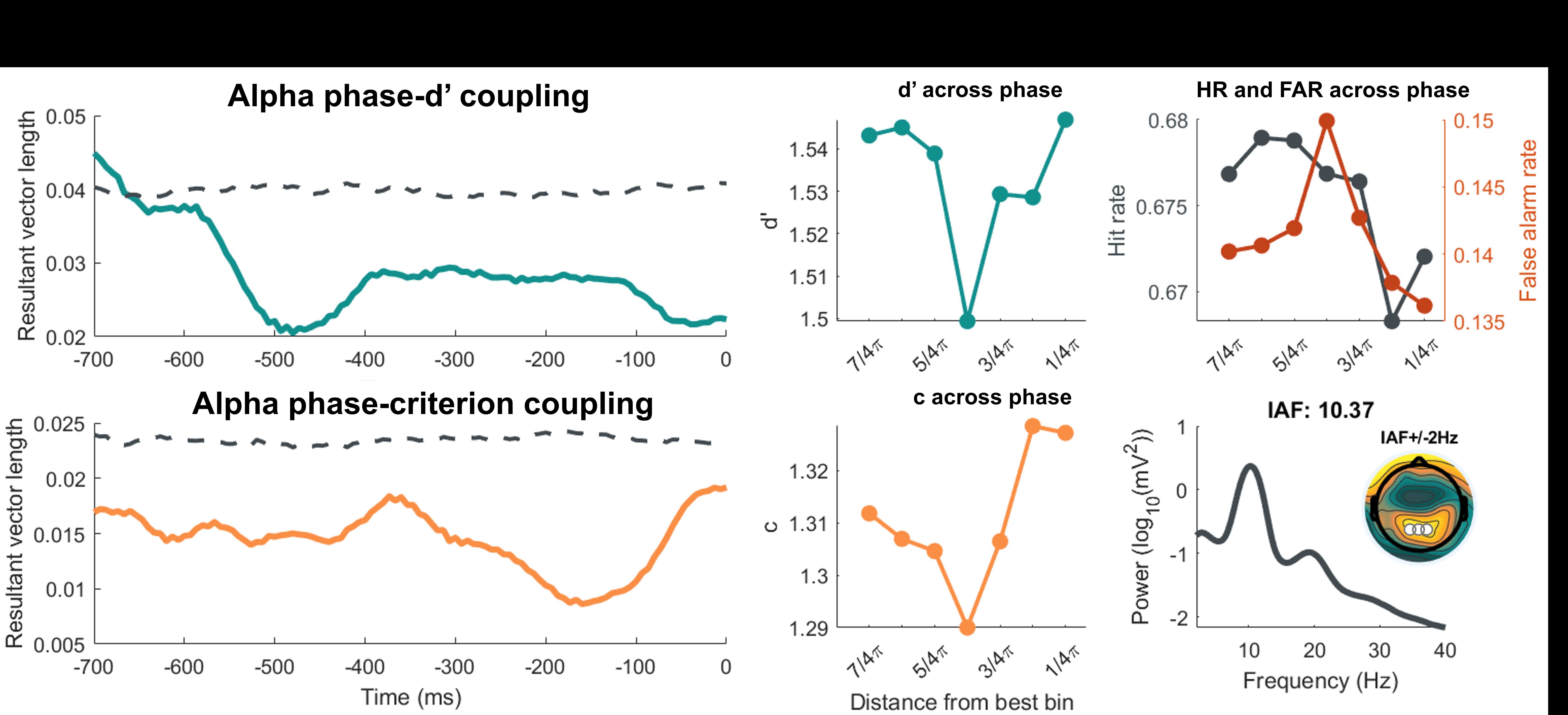

### Supplemental Figure 3

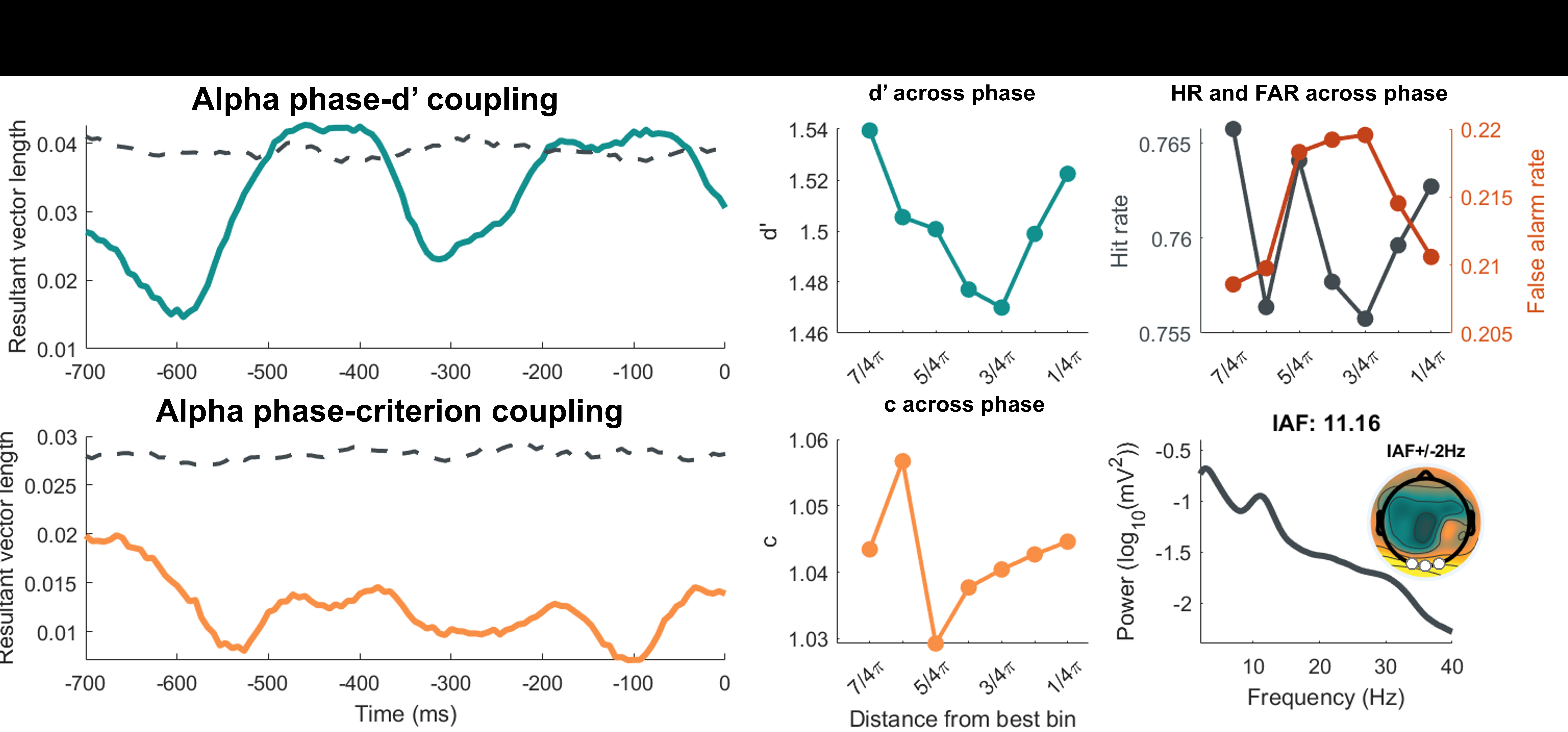

### Supplemental Figure 3

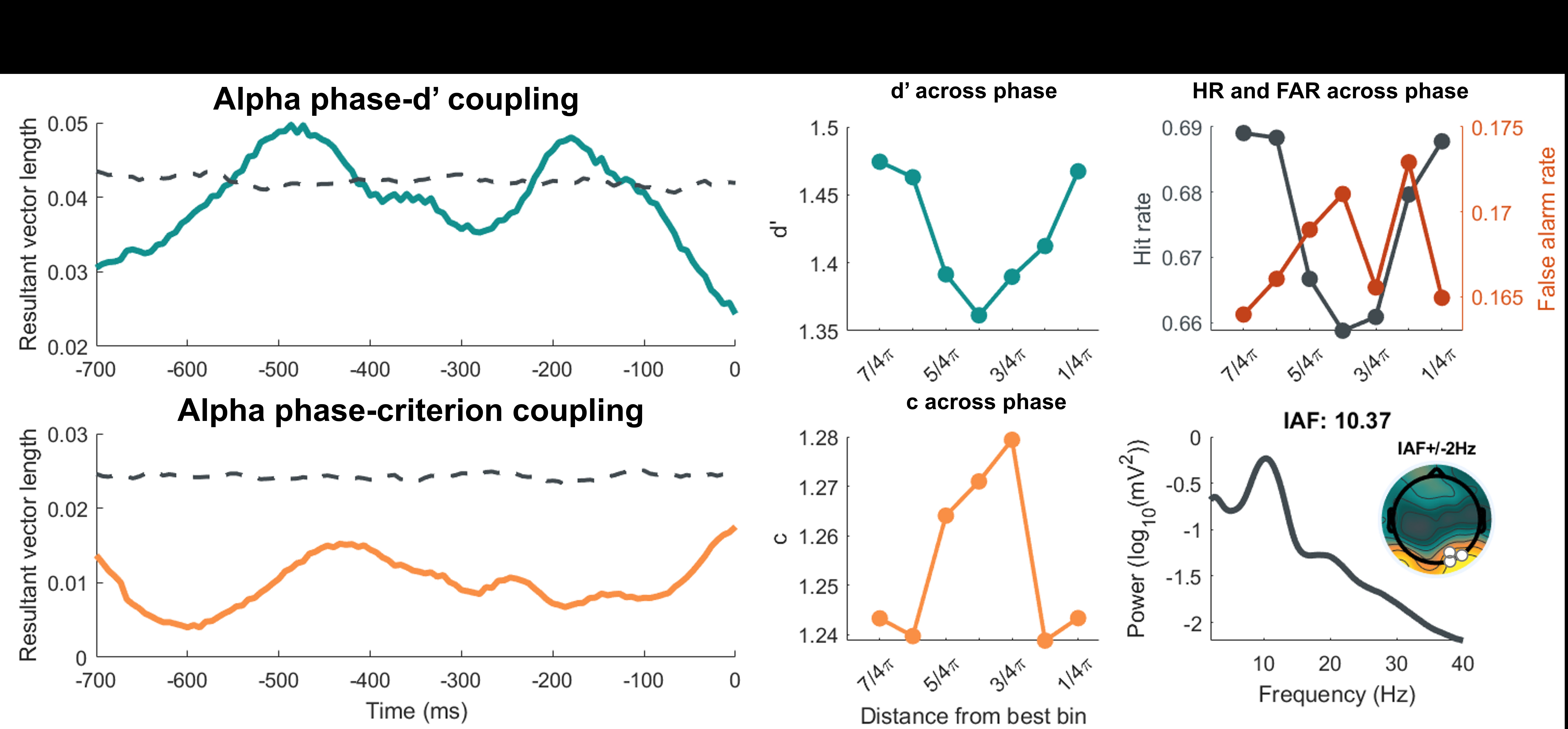

### Supplemental Figure 3

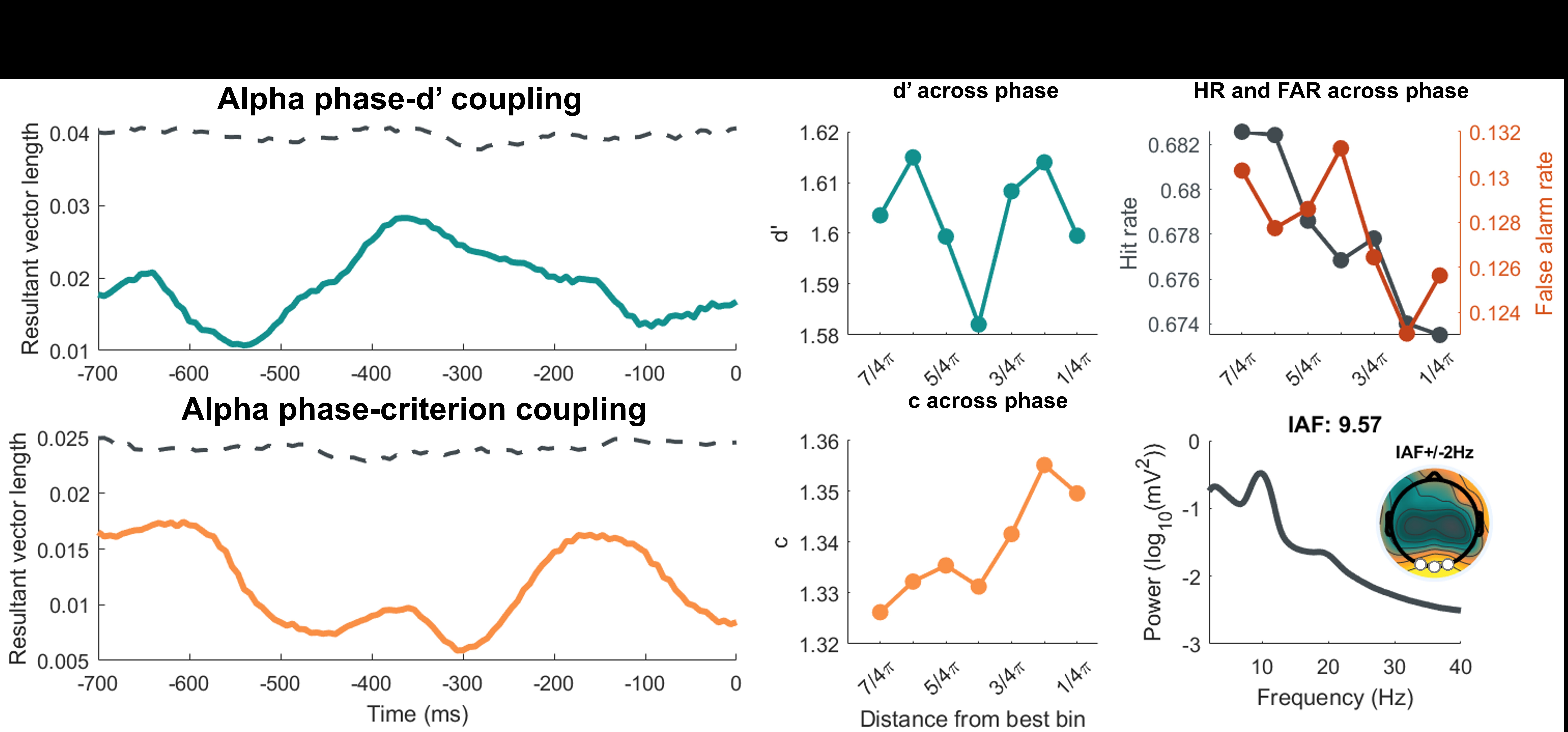
